# A tissue-level phenome-wide network map of colocalized genes and phenotypes in the UK Biobank

**DOI:** 10.1101/2021.04.30.441974

**Authors:** Ghislain Rocheleau, Iain S. Forrest, Áine Duffy, Shantanu Bafna, Amanda Dobbyn, Marie Verbanck, Hong-Hee Won, Daniel M. Jordan, Ron Do

**Author notes:** Corresponding author: Ron Do.

## Abstract

**Background:** Phenome-wide association studies conducted in electronic health record (EHR)-linked biobanks have uncovered a large number of genomic loci associated with traits and diseases. However, interpretation of the complex relationships of associated genes and phenotypes is challenging.

**Results:** We constructed a tissue-level phenome-wide network map of colocalized genes and phenotypes. First, we generated colocalized expression quantitative trait loci from 48 tissues of the Genotype-Tissue Expression project and from publicly available genome-wide association study summary statistics from the UK Biobank. We identified 9,151 colocalized genes for 1,411 phenotypes across 48 tissues. Then, we constructed a bipartite network using the colocalized signals to establish links between genes and phenotypes in each tissue. The majority of links are observed in a single tissue whereas only a few are present in all tissues. Finally, we applied the biLouvain clustering algorithm in each tissue-specific bipartite network to identify co-clusters of non-overlapping genes and phenotypes. The majority of co-clusters contains a small number of genes and phenotypes, and 88.6% of co-clusters are found in only one tissue. To demonstrate functionality of the phenome-wide map, we tested if these co-clusters were enriched with known biological and functional gene classes and observed several significant enrichments. Furthermore, we observed that tissue-specific co-clusters are enriched with reported drug side effects for the corresponding drug target genes in clinical trial data.

**Conclusions:** The phenome-wide map provides links between genes, phenotypes and tissues across a wide spectrum of biological classes and can yield biological and clinical discoveries. The phenome-wide map is publicly available at https://rstudio-connect.hpc.mssm.edu/biPheMap/.

## Background

Electronic health records (EHR)-linked biobanks coupled with genome-wide genotyping and sequencing data allows for the study of the impact of genetic variation on thousands of medical phenotypes simultaneously. Phenome-wide association analyses have been conducted in several EHR-linked biobanks and genome-wide association study (GWAS) summary statistics have been made publicly available for large biobanks such as the UK Biobank and FinnGen study [1–3]. For example, GWAS summary statistics from a phenome-wide scan of the UK Biobank (UKBB) - a prospective cohort with deep genetic and rich phenotypic data collected on approximately 500,000 middle-aged individuals (aged between 40 and 69 years old) recruited from across the United Kingdom [4] – now exists and is a rich resource in the human genetics community.

The UKBB project permits the study of the relationship of tens of thousands of genes and phenotypes simultaneously. However, a major challenge is interpretation due in large part to the complexity and heterogeneity of this data. Furthermore, there is a general lack of statistical methods available for such high-throughput analysis. Hence, only a few efforts have systematically characterized disease relationships in EHR data [5,6].

In the present study, we sought to enhance our understanding of the complex relationship of genes and phenotypes in the medical phenome by constructing a tissue-level phenome-wide network map (biPheMap) of colocalized genes and phenotypes. To construct the phenome-wide map, we first generated tens of thousands of colocalized expression quantitative trait loci (eQTL) from 48 tissues of the Genotype-Tissue Expression (GTEx) v7 project [7–9], and from ~3,800 GWAS of biological and medical phenotypes from the UKBB. We then applied a bipartite (or two-mode) network approach [10,11] followed by the biLouvain clustering method [12], to identify networks of genes and phenotypes that co-cluster together in different tissues, giving us broad insight into the biological structure of genes, phenotypes and tissues. Finally, we demonstrate functionality of the phenome-wide map by highlighting co-clusters that are biologically relevant, and by identifying enrichments of these co-clusters with both biological pathways and drug side effects in clinical trials.

## Results

We performed three steps to generate the phenome-wide network map of genes and phenotypes: 1) identification of colocalization signals of eQTLs and GWAS loci for 3,822 (before quality control) phenotypes in 48 tissues from the GTEx project v7; 2) construction of a bipartite network using the colocalization signals to establish links between genes and phenotypes in each tissue; and 3) identification of co-clusters of colocalized genes and phenotypes in each bipartite network using the biLouvain clustering algorithm. A graphical flowchart of the study is shown in **Figure 1**.

**Figure 1:**
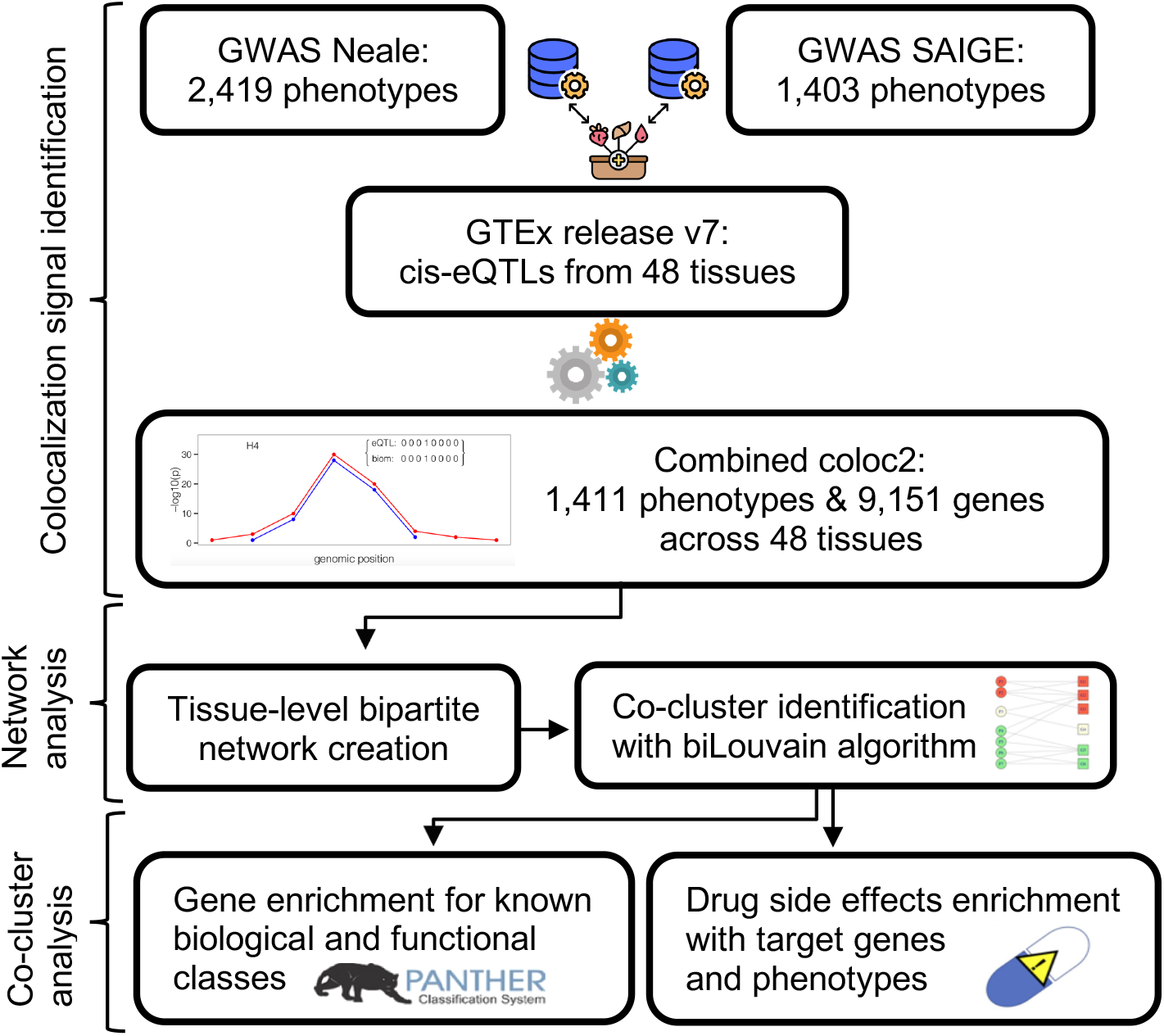
Flowchart of the study.

### Identification of colocalization signals of eQTL and GWAS loci in multiple tissues

We used coloc2 [13], along with GWAS summary association statistics for 3,822 phenotypes in UKBB [1,2] and eQTL data to identify colocalization signals in 48 tissues from the GTEx project. In total, after applying quality control (see **Methods**), we identified 9,151 unique colocalized genes for 1,411 unique phenotypes across the 48 selected tissues. Colocalization results for each tissue are reported in **Table S1, Additional file 1**. Unsurprisingly, the number of colocalized genes and phenotypes increases with respect to the GTEx tissue sample size (from n=80 for “Brain - Substantia nigra” to n=491 for “Muscle - Skeletal”), reflecting enhanced statistical power of the method to uncover colocalized genes (see **Figure S1, Additional file 2)**.

### Construction of tissue-level bipartite networks

Using the colocalized data of 9,151 genes and 1,411 phenotypes, we next created a bipartite network for each tissue. In brief, a bipartite network – also called a two-mode network – is a network in which nodes of one mode (i.e. type) are only connected to nodes of the other mode, as opposed to a unipartite (or one-mode) network commonly found in the network literature. In our colocalization results, phenotypes are not directly connected to other phenotypes, but could only be indirectly connected to each other through genes they share, while genes are indirectly connected to other genes if they appear in the same phenotype. In **Figure S2 Panel A, Additional file 2**, a typical graphical representation of a bipartite network is displayed, comprised of seven phenotypes and six genes. Associations between genes and phenotypes are indicated by links (or edges) between them.

For each tissue, **Table S2, additional file 1** displays the number of unique colocalized genes and phenotypes, along with the number of links between the two sets. When all 48 tissues are aggregated, there are 9,151 unique colocalized genes and 1,411 unique phenotypes, with 25,710 unique links between the two sets. We aggregated the tissues using an “unweighted” approach which means that if a link between a gene and a phenotype was found in more than one tissue, we counted this link only once. In fact, we observed that the majority of links between a given gene and a given phenotype are observed in one single tissue, but a few links are present in all 48 tissues (see **Figure S3**, **Additional file 2**; **Table S3, Additional file 1**).

To characterize and compare colocalization results across tissues, we computed the average degree of colocalized genes and phenotypes in each tissue. The degree of a given gene (respectively, phenotype) is simply the number of unique phenotypes (respectively, genes) connected to it in the network, the average being taken over the total number of genes (respectively, phenotypes) [14]. The average degree for both genes and phenotypes does not vary much across tissues (see **Table S2, Additional file 1**), although it increases with larger tissue sample size. When aggregating all 48 tissues, each gene is connected to an average of ~2.8 phenotypes while each phenotype is connected to an average of ~18.2 genes. The fact that the majority of gene and phenotype links are observed in a single tissue and not across all tissues, but the average degree of genes and phenotypes does not vary across tissues, suggests an architecture where tissue-specific gene regulatory mechanisms drive GWAS loci and the size and structure of these mechanisms are largely similar across different tissues.

### Identification of co-clusters in tissue-level bipartite networks

To identify structure within the phenome-wide map, we applied a newly proposed clustering algorithm, called biLouvain, which extends the well-known unipartite Louvain clustering algorithm [12]. This new algorithm efficiently identifies co-clusters of non-overlapping genes and phenotypes by maximizing a bipartite modularity measure (see **Methods** for details). **Figure S2 Panel B, Additional file 2** illustrates co-clusters identified by the biLouvain algorithm in the bipartite network of **Figure S2 Panel A, Additional file 2**.

We applied the biLouvain algorithm to identify co-clusters in each of the tissue-level bipartite networks. We identified a large number of co-clusters, ranging from 218 co-clusters for tissue “Brain - Anterior cingulate cortex (BA24)” to 314 co-clusters for “Adipose - Subcutaneous”. Across all bipartite networks, we observed that the majority of co-clusters had a small number of genes and phenotypes, on one hand, whereas a few co-clusters had a large number of genes and phenotypes, on the other hand (**Figure 2, Panels A-B**). Across the 48 tissues, the vast majority of co-clusters (8,389 / 9,472 = 88.6%) were found in only one tissue (**Figure 2, Panel C**). Hence, the structure of the phenome-wide map involves hundreds of isolated tissue-specific subnetworks comprised of a small number of interrelated genes and phenotypes. Large co-clusters were also identified, although these are the exception rather than the norm. The complete list of genes and phenotypes per co-cluster in each tissue is provided in **Table S4, Additional file 1**.

**Figure 2:**
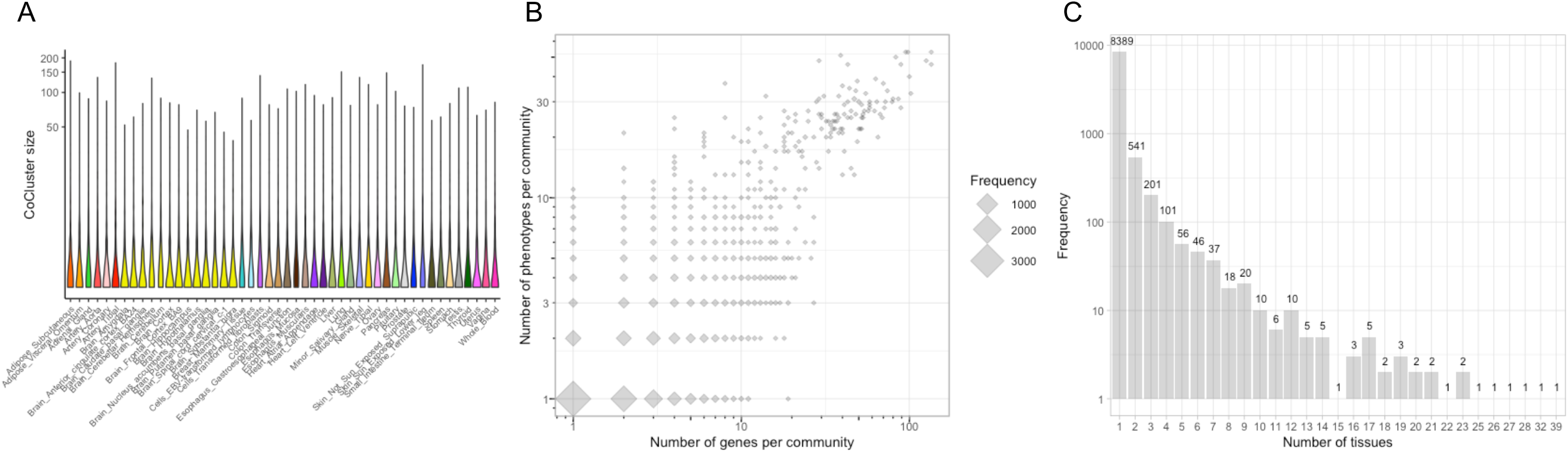
Characteristics of biLouvain co-clusters across tissues. A) Violin plot of co-cluster size (number of genes + number of phenotypes) for each tissue. Y-axis is displayed in log scale; B) Number of genes and phenotypes per co-cluster identified by the biLouvain algorithm. Diamonds are proportional to frequency of co-cluster size across all 48 tissues. Both axes are displayed in log scale; C) Number of unique co-clusters and how many times they appear in a given number of tissues. Y-axis is displayed in log scale.

### Enrichment analysis of co-clusters with biological and functional gene classes

To demonstrate functionality of the phenome-wide map, we tested if the identified biLouvain co-clusters were enriched with known biological and functional gene classes. We selected 183 co-clusters consisting of 10 genes or more, and performed enrichment analysis using PANTHER [15,16] on four different annotation types: Biological process (2,064 gene ontology (GO) terms), Cellular component (520 GO terms), Molecular function (532 GO terms), and 164 different Pathways. For each co-cluster and each annotation type, we selected the minimal p-value of all Fisher “overrepresentation” tests, and plotted it against the expected minimal p-value under the null hypothesis of no enrichment (see **Methods** for details). All four annotation types demonstrated significant enrichment. We observed enrichment in GO terms related to i) antibody-mediated immune response, upregulation response to biotic stimulus, glutathione metabolism, zymogen activation, downregulation of blood pressure, and cellular nitrogen compound metabolism in seven co-clusters in the Biological process annotation; ii) outer surface of cytoplasmic membrane, and obsolete intracellular part in two co-clusters in the Cellular component annotation; iii) signaling receptor binding, metallopeptidase activity, zinc ion binding, NADH-dependent glyoxylate reductase, and phosphatase activity in five co-clusters in the Molecular function annotation; and iv) toll-like receptor signaling pathway, muscarinic acetylcholine receptor 2 and 4 signaling pathway, serine and glycine biosynthesis, and heterotrimeric G-protein signaling pathway-Gq alpha and Go alpha mediated pathway in six co-clusters in the Pathway annotation.

As an example, the most significant pathway detected by PANTHER is a toll like receptor signaling pathway for co-cluster 111 comprised of hypothyroidism/myxoedema and levothyroxine sodium medication, and genes *TLR1*, *TLR6* and *TLR10* in the “Cells - EBV-transformed lymphocytes” tissue (**Figure 3**; **Table 1**; **Figure S4 Panel A, Additional file 2**). Toll like receptor 1 (*TLR1*), 6 (*TLR6*) and 10 (*TLR10*) genes are located in the same gene cluster on chromosome 4p14, and they play a fundamental role in pathogen recognition and activation of innate immunity [17]. Previous studies have shown that *TLR1* and *TLR10* are linked to Graves’ disease [18] and Hashimoto’s disease [19], which are clinical subtypes of autoimmune thyroid diseases. Furthermore, variants in *CD226* [20], and *RASGRP1* [21] were found to be associated with autoimmune thyroid diseases and with thyroid preparations (H03A medication class, which comprises levothyroxine sodium) [22].

**Table 1:**
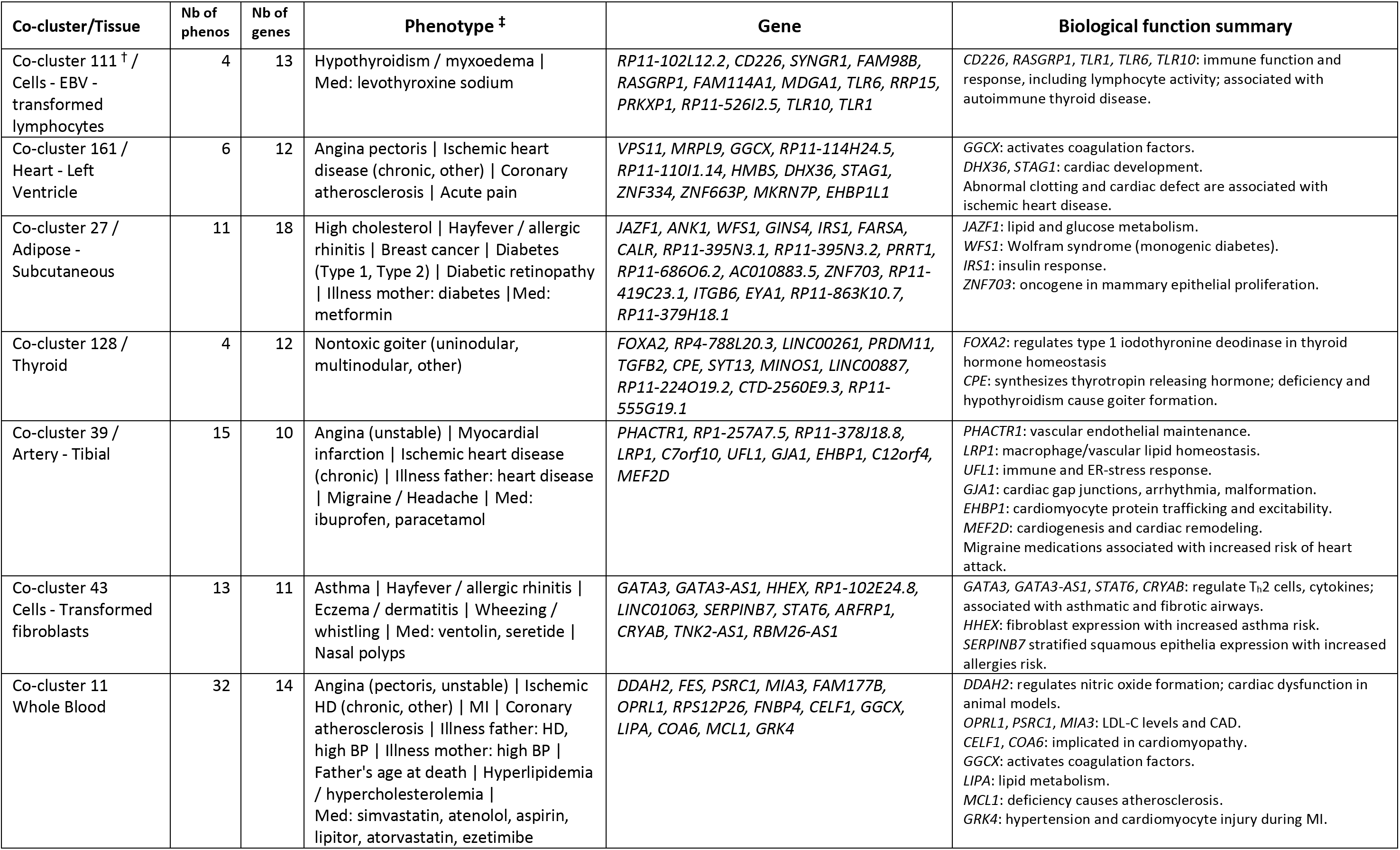
Selection of interesting co-clusters in relevant tissues. Nb: Number; HD: Heart disease; MI: Myocardial infarction; BP: Blood pressure; Med: Medication; LDL-C: low-density lipoprotein cholesterol; CAD: coronary artery disease. † Co-cluster most significant in PANTHER Pathway annotation. ‡ For the sake of simplification, we grouped phenotypes with similar names in UKBB and/or matching ICD-10 codes and PheCodes.

**Figure 3:**
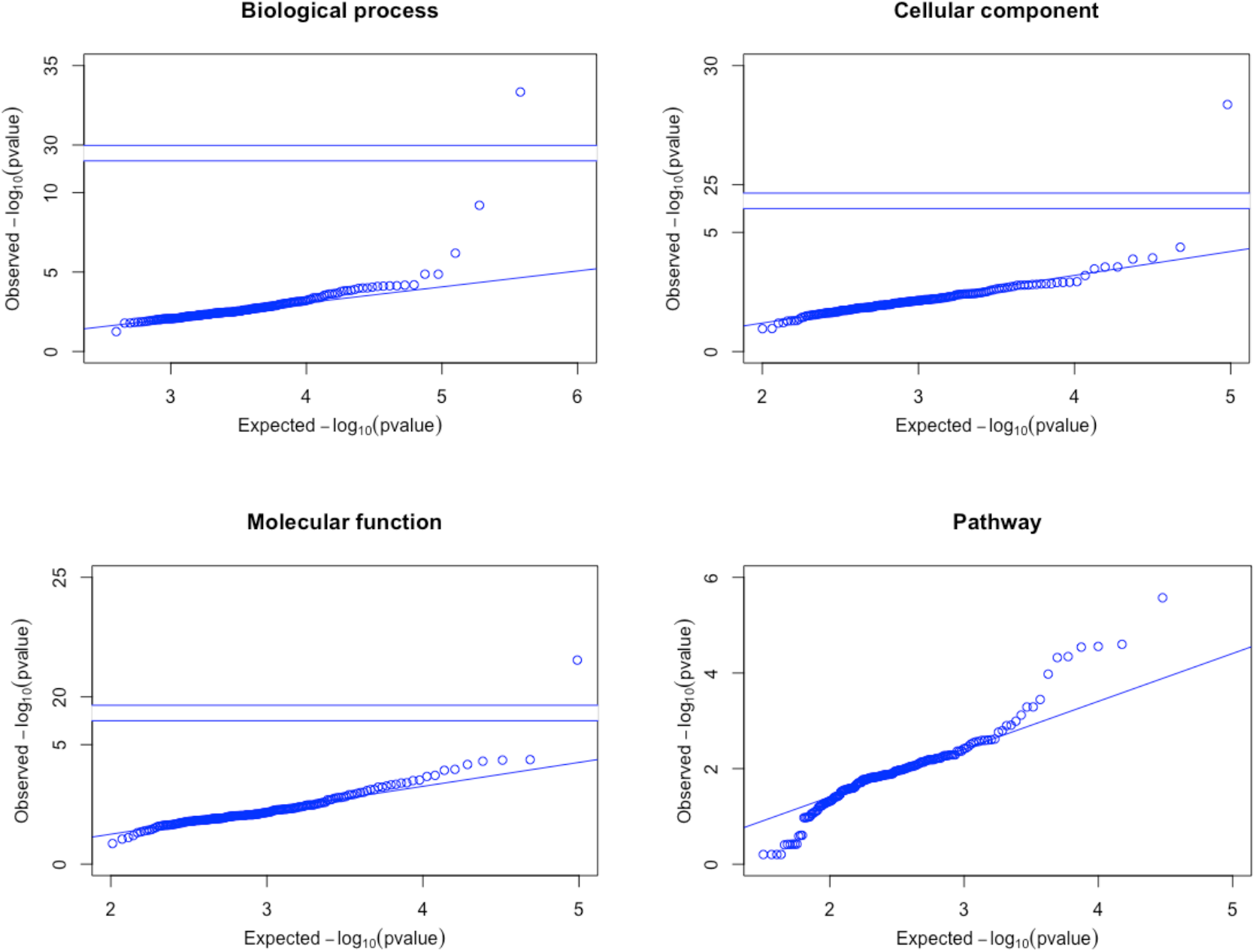
PANTHER enrichment analysis of selected biLouvain co-clusters across tissues. In each annotation type, the minimum p-value across all GO terms is displayed for all 183 co-clusters selected. Some plots show a breakdown in the y-axis to help display very small p-values.

In addition to the PANTHER gene set enrichment analysis, we identified seven co-clusters comprised of known relationships between genes and phenotypes in relevant tissues (**Table 1**), providing strong biological relevance. For example, the same gene *GGCX* appears in two co-clusters in related tissues, co-cluster 161 in “Heart - Left Ventricle” and co-cluster 11 in “Whole Blood” (**Table 1**; **Figure S4 Panels B-C, Additional file 2**). Gamma-glutamyl carboxylase (*GGCX*) encodes an integral membrane protein of the rough endoplasmic reticulum that carboxylates glutamate residues of vitamin K-dependent proteins to gamma carboxyl glutamate. Vitamin K-dependent proteins affect a number of physiologic processes including blood coagulation, inflammation, and prevention of vascular calcification [23]. Furthermore, a meta-analysis including the UKBB data identified an intronic variant in *GGCX* associated with coronary artery disease (CAD), with inclusion or exclusion of angina [24]. Taken together, these results suggest that our newly identified co-clusters contain relevant biological information and provide both novel and known functional links between genes and phenotypes.

### Drug side effect enrichment in co-clusters

To further demonstrate functionality of the phenome-wide map, we tested for enrichment of colocalized genes and phenotypes in the phenome-wide map with drug side effects in corresponding drug target genes in an adapted integrated clinical trial dataset of 1,780 drugs [25] (see **Methods**) in each of the 48 GTEx tissues, adjusting for co-cluster grouping. In almost all tissues, we observed marked enrichments of reported drug side effects with colocalized target genes and phenotypes (**Figure 4**; **Table S5, Additional file 1**), with the strongest associated odds ratio (OR) observed in “Minor Salivary Gland” (OR = 86.43, 95% confidence interval (CI) = (11.596 – 11,078.85), p-value = 6.12 × 10^−11^), in “Breast - Mammary Tissue” (OR = 10.98, 95% CI = (3.514 – 54.897), p-value = 2.19 × 10^−6^) and in “Cells - EBV-transformed lymphocytes” (OR = 7.03, 95% CI = (2.070 - 28.367), p-value = 1.43 × 10^−3^). Nearly half of the tissues (20 out of 48) show very significant enrichment with p-value < 2.2 × 10^−16^ (**Table S5, Additional file 1**). These results suggest that the phenome-wide map can inform the drug safety profiles of candidate drug therapeutics.

**Figure 4:**
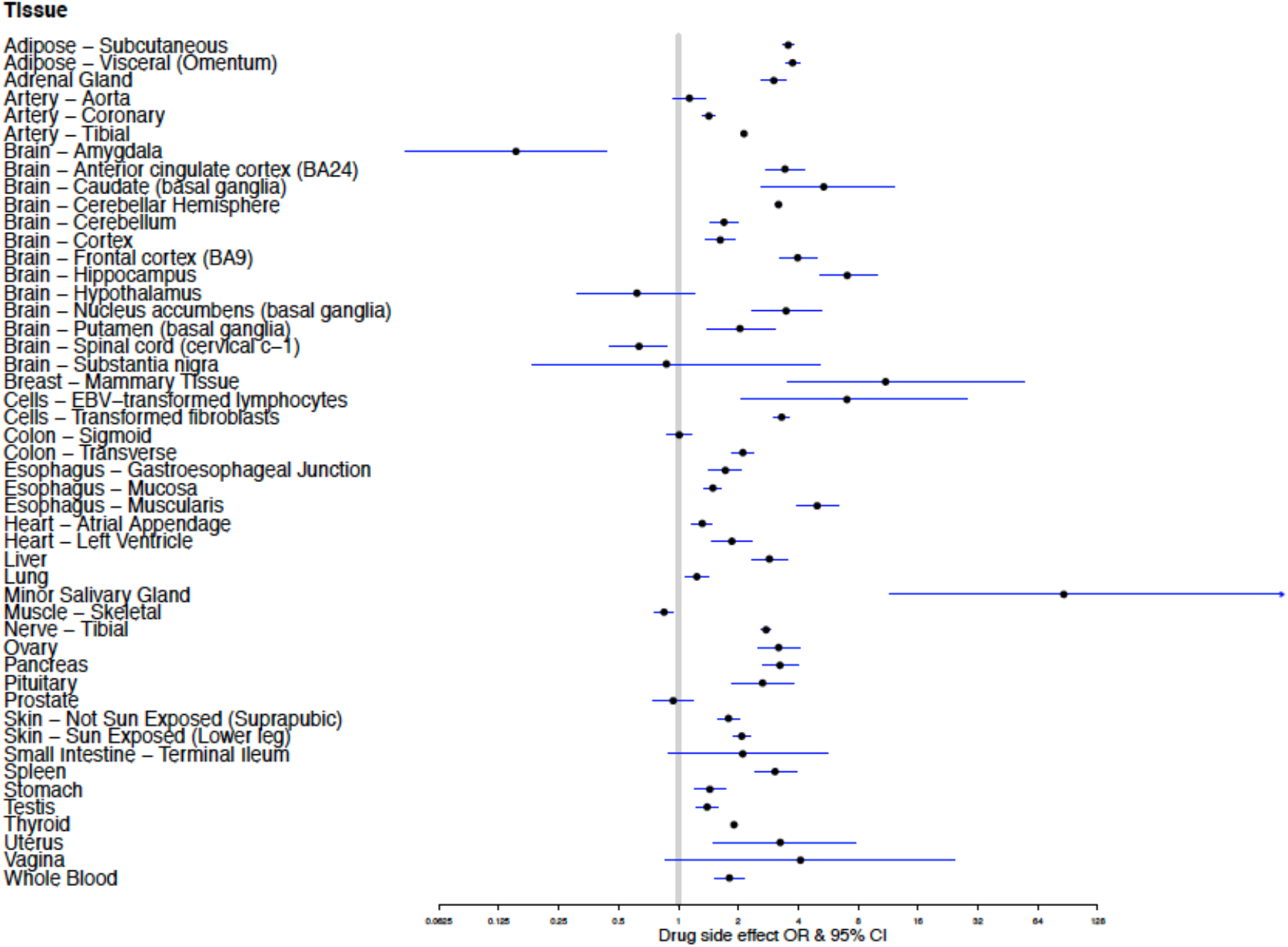
Drug side effect enrichment with colocalized target genes and phenotypes in each tissue. The odds ratio (OR) and its corresponding 95% confidence interval (CI) of drug side effect enrichment is depicted for each tissue. X-axis is displayed in log-scale.

## Discussion

In the present study, we have constructed a tissue-level phenome-wide network map, called biPheMap, of colocalized genes and phenotypes using a bipartite network and biLouvain clustering approach on 1,411 phenotypes and eQTL data from 48 tissues from the GTEx project. In the phenome-wide map, we observed the following: 1) the majority of colocalized gene and phenotype links are observed in a single tissue, implying that tissue-specific gene regulatory mechanisms drives phenotypic variation; 2) the majority of co-clusters are comprised of a small number of gene and phenotype links; 3) specific co-clusters are enriched with functional gene set annotations; 4) specific co-clusters are identified with biologically relevant gene, phenotype and tissue functions; 5) tissue-level co-clusters are enriched with reported drug side effects for the corresponding drug target genes.

While most network analyses have focused on unipartite networks, the present study used the less familiar bipartite approach. Many such bipartite networks have been studied in different contexts: actor-movie network in cinema industry, author-scientific paper networks in academia, pollinator-plant in ecological networks, etc., but their topological features and related metrics are unique and different from their more classical unipartite counterpart. A simpler analysis could have been proposed by “projecting” the bipartite network into two unipartite networks to produce a gene-gene network and a phenotype-phenotype network. However, this projection method entails a loss of information since the original links between genes and phenotypes are no longer available. Such a projection approach was employed in Verma et al. [26] to create a “disease-disease” network where more than 500 diagnosis codes were linked on the basis of shared variant associations.

An important feature of the phenome-wide map is the exploration and discovery of co-clusters of related genes and phenotypes. So far, few community detection algorithms in bipartite networks could be run in a reasonable amount of time. One fast and precise algorithm is the biLouvain algorithm [12] which maximizes bipartite modularity, an extension of the modularity measure found in unipartite network clustering algorithms. **Table 1** displays various examples of gene-phenotype co-clusters confirming known genetic associations and also suggesting unsuspected etiological links between phenotypes. For example, many epidemiological and genetic studies have suggested shared loci between migraine and CAD, and one study identified gene *PHACTR1* as the strongest shared locus between the two disorders [27]. However, some co-clusters might also consist of phenotypes being observed as a consequence of another phenotype. For example, we observed lipid-lowering medications in the same co-cluster as lipid disorders.

There are some limitations to our study that deserve mention. First, some phenotypes are highly correlated in UKBB, and therefore, colocalized signals were sometimes redundant in our phenome-wide map. Second, we applied stringent quality control in our phenotype selection to avoid reporting false colocalized loci. This came at the expense of missing some loci, especially if the leading associated variant in a locus is rare (minor allele frequency < 0.1%). Third, the literature cited to support the links between genes and phenotypes of co-clusters displayed in **Table 1** relies heavily on genetic associations found in the GWAS summary statistics in UKBB, which is comprised solely of British white individuals. Incorporation of findings from diverse ancestry populations will be necessary to yield more genetic association phenotypes. Finally, we used the bipartite network approach over the more common unipartite approach, which limited the set of tools to analyze our results. Fortunately, the bipartite network and its characteristics is gaining more attention in the network literature, and future methodological developments will expand the range of tools and analyses that could be performed in this type of networks.

## Conclusions

In conclusion, we showed that the phenome-wide map can be a useful resource to understand gene, phenotype and tissue links across a wide spectrum of biological classes and diseases. We expect that further interrogation of the phenome-wide map will yield more biological and clinical discoveries.

## Methods

### Datasets

The present study uses two resources: a) the UK Biobank (UKBB) project; and b) the Genotype-Tissue Expression (GTEx) project. The UKBB project is a prospective EHR-linked cohort with deep genetic and rich phenotypic data collected on approximately 500,000 middle-aged individuals (aged between 40 and 69 years old) recruited from across the United Kingdom [4]. The GTEx project is a resource database and associated multi-tissue bank aimed at studying the relationship between genetic variation and gene expression in different human tissues [7–9].

### Colocalization method

We integrated multiple association datasets to assess whether two association signals, one from a genome-wide association study (GWAS) on a phenotype, and the other from expression quantitative trait locus (eQTL) analysis in a tissue, overlap in such a matter that they are consistent with a shared causal gene. This approach, referred to as colocalization, was conducted using coloc2 [28], an enhancement of the previously published method coloc [13]. The coloc2 method is a Bayesian approach which computes the posterior probability that a genetic variant is both associated with the phenotype and the gene expression level in the tissue (denoted PPH4 in [13]). We defined a colocalized signal using PPH4 ≥ 0.80, as described previously [13].

### GWAS and eQTL summary statistics

coloc2 requires both GWAS summary data and eQTL association summary data. For GWAS data, we used two sets from the UKBB project. The first set of results are GWAS association test statistics publicly available from the Neale lab (Round 1 in 2,419 phenotypes). We selected variants with minor allele frequency (MAF) > 0.1% and with association p-value < 5×10^−5^. More details on the data quality control and the full list of phenotypes can be found at [1]. We further used a second set of UKBB GWAS association statistics computed by the SAIGE testing method [29]. In total, 1,403 case-control phenotypes (PheCodes) were available. We selected variants with MAF > 0.1% and with association p-value < 5×10^−3^. Full datasets and list of PheCodes can be downloaded at [2]. For eQTL association signals, we used data from Analysis V7 of the GTEx project [30]. We restricted our study to the list of 48 tissues (from 620 donors) having a sample size of at least 80. eQTLs with MAF > 1% were considered as input for coloc2. Statistically significant cis-eQTLs were selected as detailed in [31] and available on the GTEx Portal.

After running coloc2 on the Neale GWAS data, we performed stringent quality control. First, we removed results from phenotypes related to cause of death (since these phenotypes generally had very low number of cases), and we also removed case-control phenotypes with less than 1,250 cases (or controls), except phenotypes showing prior gene/locus association as reported in the NHGRI-EBI GWAS catalog [32,33]. The rationale for excluding phenotypes with less than 1,250 cases (or controls) is based on the recommendation by Neale to keep only variants with at least 25 minor alleles in the sample of cases (or controls), in order to avoid inflation in association test statistics due to extreme case-control ratio imbalance and ensuring reliable p-value computation [34]. With the Neale GWAS data, we retained coloc2 results for 496 continuous and binary phenotypes. In the same vein, after running coloc2 with SAIGE association data, we excluded case-control phenotypes with less than 200 cases, as recommended by the authors [29]. With SAIGE data, we generated coloc2 results for 915 case-control (PheCode) phenotypes.

### Construction and descriptive statistics of bipartite networks

To construct the phenome-wide map of genes and phenotypes using the colocalized data of 9,151 genes and 1,411 phenotypes, we created a bipartite network for each tissue. In brief, a bipartite network, also called a two-mode network, is a network in which nodes of one mode (i.e. type) are only connected to nodes of the other mode (for a review, see [10,11]). Associations between phenotypes and genes are indicated by links or edges between them.

To characterize and compare colocalization results across tissues, we computed descriptive statistics adapted to bipartite networks. We computed the average degree of colocalized genes and phenotypes in each tissue. The degree of a given gene (respectively, phenotype) is simply the number of unique phenotypes (respectively, genes) connected to it in the network, the average being taken over the total number of genes (respectively, phenotypes) [14].

### biLouvain clustering algorithm

For each tissue, it is expected that the bipartite network of coloc2 results will tend to “cluster” in small groups of related phenotypes with their causally associated genes. To uncover clustering within each network, we applied a newly proposed clustering algorithm, called biLouvain, which extends the well-known unipartite Louvain clustering algorithm. This new algorithm efficiently identifies co-clusters, also called communities, of non-overlapping genes and phenotypes by maximizing a bipartite modularity measure (see [12] for details).

### PANTHER enrichment analysis

To test if biLouvain co-clusters were enriched with some functional gene classes, we selected 183 co-clusters consisting of 10 genes or more, and input them into the online PANTHER enrichment analysis tools [15,16,35]. We applied Fisher “overrepresentation” tests on four different annotation types: Biological process (2,064 Gene Ontology (GO) terms), Cellular component (520 GO terms), Molecular function (532 GO terms), and 164 different Pathways. For each co-cluster and each annotation type, we took the minimal p-value of all Fisher tests, and plotted it against the expected minimal p-value under the null hypothesis of no enrichment. We assumed that p-values within each annotation type are independently distributed uniformly over the interval (0,1), which represents a conservative approach. Note that the minimal p-value of *n* independent p-values from a Uniform(0,1) is not uniformly distributed under the null: its cumulative density function is instead given by Prob(*X* ≤ *x*) = 1 − (1 − *x*)^*n*^, 0 < *x* < 1. In each panel of **Figure 3**, we plotted a straight line with slope equal to 1, which crosses the y-axis at x = (observed 1^st^ quartile – expected 1^st^ quartile) using the above expected cumulative density function. Gene enrichment was deemed significant if the minimal p-value was less than 0.05/(183 × 4) = 6.8 × 10^−5^.

### Drug side effects enrichment

To test if the identified co-clusters were enriched with target genes with reported drug side effects for relevant phenotypes, we used our integrated clinical-genetic drug side effect dataset described and published in [25]. This consists of 1,780 drugs and their gene targets with side effect and coloc2 results mapped to 48 GTEx terms (see [25] for mapping approach). To combine this dataset with our biLouvain co-clusters, we recorded the side effect, colocalized phenotype and tissue for each drug-gene pair (rather than collapse all gene targets per drug as in [25]). We restricted the dataset to drug genes present in each tissue-level network by mapping these genes to their corresponding co-cluster for that tissue. To evaluate the association between colocalized phenotype and drug side effect for each tissue-level network, we selected genes that were also colocalized in the same tissue as in our integrated clinical-genetic drug side effect dataset. For each tissue, we used a Firth logistic regression model (R package logistf) with side effect as the outcome variable and phenotype as the predictor, adjusting for the co-cluster grouping.

## Declarations

### Ethics approval and consent to participate

Not applicable

### Consent for publication

Not applicable

### Availability of data and materials

All colocalized results analyzed in this study are available through a R Shiny app called biPheMap at https://rstudio-connect.hpc.mssm.edu/biPheMap/.

This research has been conducted using the UK Biobank Resource under Application Number ‘16218’. UK Biobank data is available to researchers upon approval of an application form at https://www.ukbiobank.ac.uk/. The GTEx Analysis V7 dataset can be freely downloaded at https://www.gtexportal.org/home/datasets. At the time of our colocalization analysis, we utilized Round 1 of Benjamin Neale’s lab GWAS summary statistics in the UK Biobank. The Round 1 results are no longer accessible and has since been replaced by a more recent Round 2 which can be freely downloaded at http://www.nealelab.is/uk-biobank. Seunggeun Lee’s lab GWAS summary statistics in the UK Biobank using SAIGE can be freely downloaded at https://www.leelabsg.org/resources.

R codes to run coloc2 are available at https://github.com/Stahl-Lab-MSSM. The biLouvain algorithm, described in [12], may be downloaded and installed by following instructions at https://github.com/paolapesantez/biLouvain.

The PANTHER classification system is available at http://www.pantherdb.org/.

### Authors’ contributions

GR and RD conceived and planned the study. HHW managed and created the pipeline to generate colocalization results. GR, DMJ, HHW, ISF, AiD, AmD, MV, RD analyzed and/or interpreted the colocalization results. DMJ performed the PANTHER enrichment analysis. AiD performed the drug side effect enrichment analysis. SB implemented and deployed the biPheMap R shiny app. GR created all figures and tables, with assistance from ISF, AiD, SB. GR and RD drafted and revised the manuscript. All authors read and approved the final manuscript.

## Acknowledgments

We thank Dr. Benjamin Neale’s and Dr. Seunggeun Lee’s group for generously sharing GWAS summary statistics in the UK Biobank. We would also like to thank the donors and their families for making organ and tissue donations to the GTEx project.

## Supplementary information

**Additional file 1 (XLSX) - Tables S1-S5**

**Table S1**. **Colocalization results across the 48 selected tissues in GTEx (9,151 unique colocalized genes for 1,411 unique phenotypes)**.

Each row represents a colocalized signal between the gene and the phenotype in the corresponding tissue. In the Phenotype column, the prefix “N_” stands for Neale’s GWAS association statistics, while “S_” stands for SAIGE GWAS association statistics.

**Table S2**. **Descriptive statistics of tissue-level bipartite networks**. For each bipartite network, the number of phenotypes, genes and links are reported, along with the average degree across all phenotypes and all genes, respectively.

**Table S3**. **Colocalization results aggregated over tissues**.

Each row represents a unique colocalized signal between the gene and the phenotype, and reports in how many tissues this signal is found, along with the list of these tissues. In the Phenotype column, the prefix “N_” stands for Neale’s GWAS association statistics, while “S_” stands for SAIGE GWAS association statistics.

**Table S4**. **List of all co-clusters found by the biLouvain algorithm for each tissue-level bipartite network**.

The number of phenotypes and genes in each co-cluster are provided in the last two columns (number of phenotypes and number of genes). In the Phenotype column, the prefix “N_” stands for Neale’s GWAS association statistics, while “S_” stands for SAIGE GWAS association statistics.

**Table S5**. **Drug side effect enrichment with colocalized target genes and phenotypes in each tissue, adjusted for co-cluster grouping**.

The odds ratio (OR) and its corresponding 95% confidence interval (CI) of drug side effect enrichment is displayed for each tissue, along with the p-value for enrichment.

**Additional file 2 (DOCX) - Figures S1-S4**

