## Additional file 2 for "A tissue-level phenome-wide network map of colocalized genes and phenotypes in the UK Biobank"

**Figure S1**. **Bubble plot of the number of colocalization signals for each tissue.** Number of genes and number of phenotypes are given for each tissue, and the size of the bubble is proportional to the tissue sample size in the GTEx project.

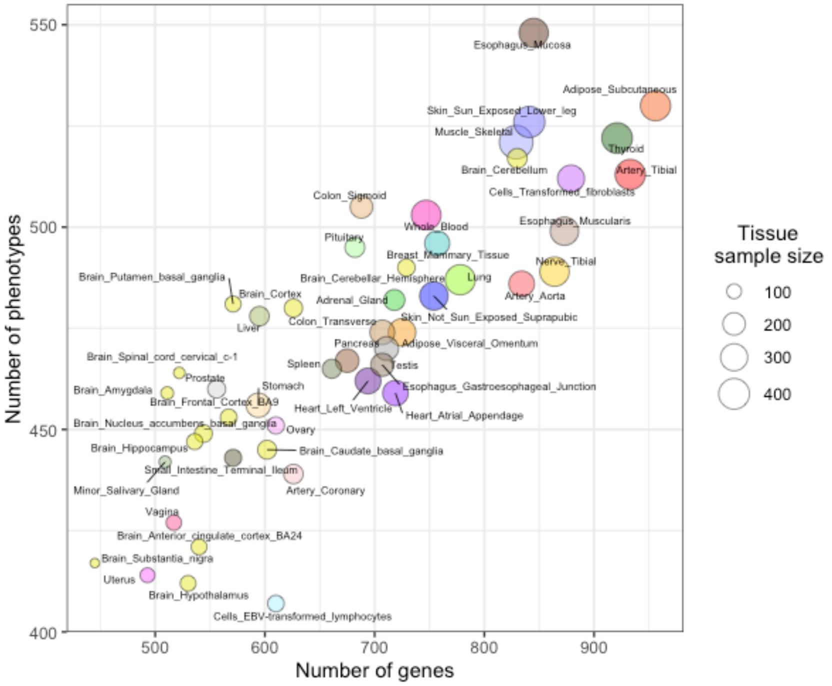

**Figure S2**. **Graphical representation of a bipartite graph.**
**Panel A**: Bipartite graph with 7 phenotypes (P1 to P7) and 6 genes (G1 to G6).
**Panel B**: Algorithm biLouvain identifies 3 co-clusters made of phenotypes and genes: {P1,P2,G1,G2,G3} in red, {P3,G4} in yellow, and {P4,P5,P6,P7,G5,G6} in green.

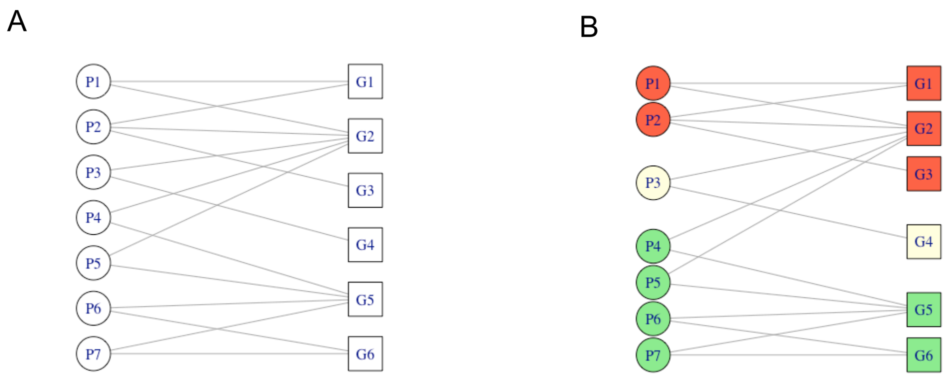

**Figure S3**. **Number of gene-phenotype links observed in number of tissues.** Distribution of the number of times a link (phenotype-gene pair) appears across all tissues (varies from 1 to 48). Both axes are displayed in log scale.

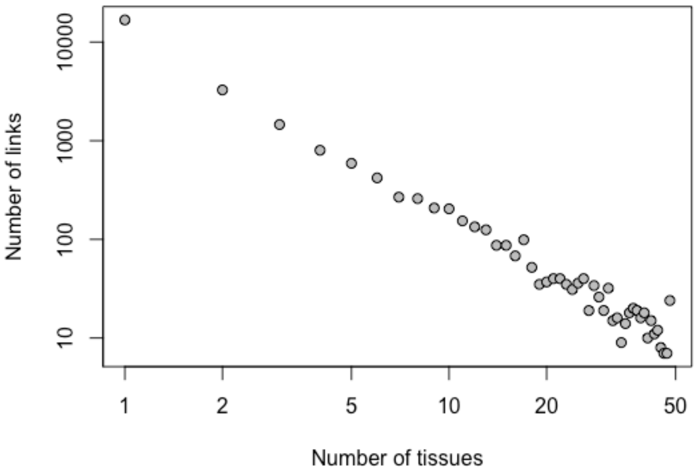

**Figure S4**. **Selected plots of subgraph induced by co-clusters.**

A) Co-cluster 111 in tissue “Cells - EBV-transformed lymphocytes”

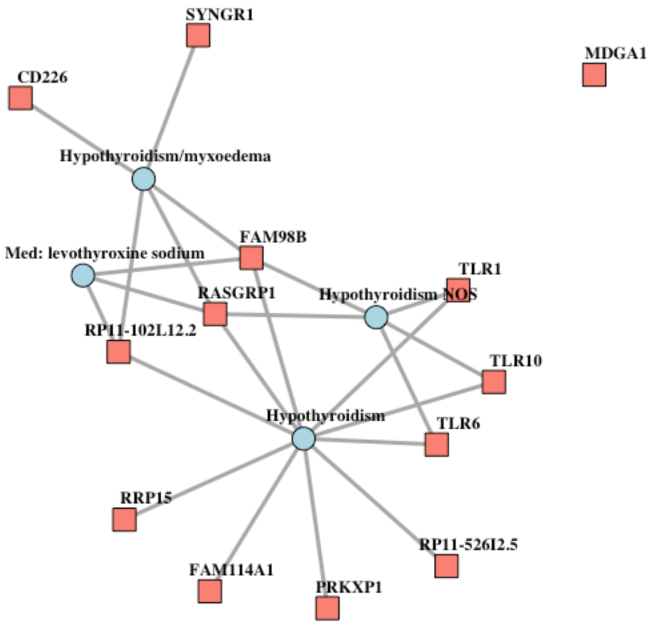


B) Co-cluster 161 in tissue “Heart - Left Ventricle”

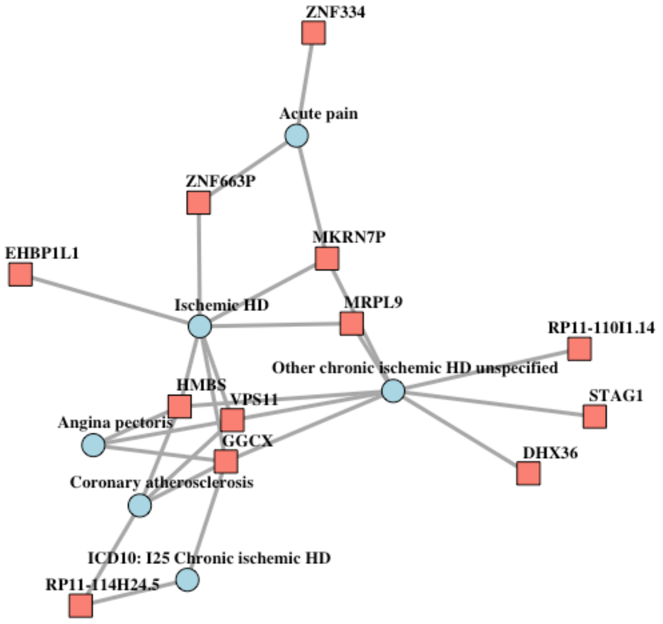


HD: heart disease

C) Co-cluster 11 in tissue “Whole blood”

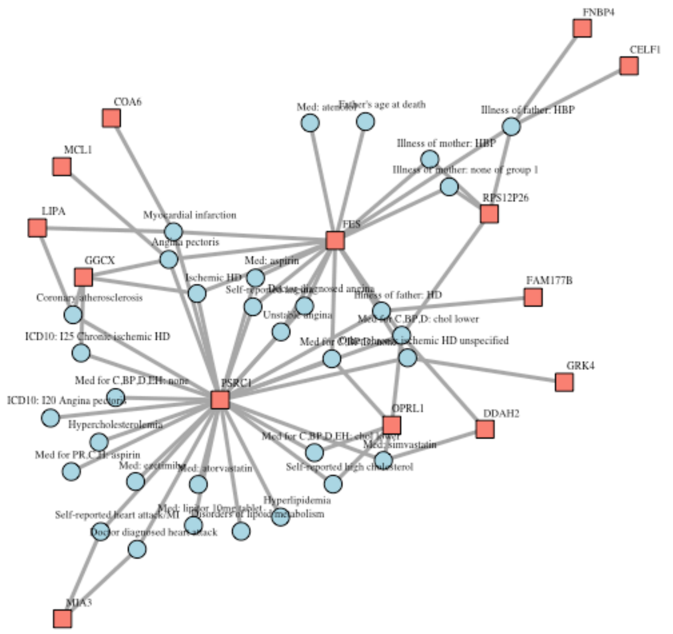


Illness of mother group 1 refers to UKBB data-field 20110 which comprises heart disease, stroke, high blood pressure, chronic bronchitis/emphysema, Alzheimer's disease/dementia, diabetes;
Med for C,BP,D: Medication for cholesterol, blood pressure, diabetes;
Med for C,BP,D, EH: Medication for cholesterol, blood pressure, diabetes, or take exogenous hormones;
Med for PR,C,H: Medication for pain relief, constipation, heartburn;
HD: heart disease;
MI: myocardial infarction;
HBP: high blood pressure;
chol lower: cholesterol lowering
